# Behavioral and cortical arousal from sleep, muscimol-induced coma, and anesthesia by direct optogenetic stimulation of cortical neurons

**DOI:** 10.1101/2024.04.19.590330

**Authors:** Rong Mao, Matias Lorenzo Cavelli, Graham Findlay, Kort Driessen, Michael J Peterson, William Marshall, Giulio Tononi, Chiara Cirelli

## Abstract

The cerebral cortex is widely considered part of the neural substrate of consciousness. However, while several studies have demonstrated that stimulation of subcortical nuclei can produce EEG activation and restore consciousness, so far no direct causal evidence has been available for the cortex itself. Here we tested in mice whether optogenetic activation of cortical neurons in posterior parietal cortex (PtA) or medial prefrontal cortex (mPFC) is sufficient for arousal from three behavioral states characterized by progressively deeper unresponsiveness: sleep, a coma-like state induced by muscimol injection in the midbrain, and deep sevoflurane-dexmedetomidine anesthesia. We find that cortical stimulation always awakens the mice from both NREM sleep and REM sleep, with PtA requiring weaker/shorter light pulses than mPFC. Moreover, in most cases light pulses produce both cortical activation (decrease in low frequencies) and behavioral arousal (recovery of the righting reflex) from brainstem coma, as well as cortical activation from anesthesia. These findings provide evidence that direct activation of cortical neurons is sufficient for behavioral and/or cortical arousal from sleep, brainstem coma, and anesthesia.

## Introduction

There is a long tradition of trying to wake up subjects from sleep or anesthesia by stimulating different parts of the brain, starting from the identification of the brainstem activating system by Moruzzi and Magoun (Moruzzi and Magoun, 1949). In that seminal study, electrical stimulation of the reticular formation rapidly converted the synchronized, high voltage low frequency EEG pattern induced by chloralose anesthesia into an “activated” pattern with low voltage fast activity (Moruzzi and Magoun, 1949). EEG activation occurred also after direct stimulation of the intralaminar thalamus, but it was still achievable after this area was lesioned (Moruzzi and Magoun, 1949). Thus, this early result suggested that certain thalamic nuclei may be dispensable for the EEG activating response, even though many excitatory projections from the reticular activating system reach the cortex via the thalamus.

Since then (Moruzzi and Magoun, 1949), the view of the activating system has evolved from a monolithic reticular core to an ensemble of distinct cell groups that promote arousal, including cholinergic, noradrenergic, dopaminergic, and glutamatergic neurons. These cell groups have diffuse projections to the cerebral cortex and thalamus and share the property of being, on average, more active during waking than during non-rapid eye movement (NREM) sleep, when the EEG is dominated by synchronous, high voltage slow waves (Brown et al., 2012; Scammell et al., 2017). They also have descending projections to the caudal brainstem and spinal cord, whose effects on muscle tone and behavioral arousal were not studied in early experiments (Moruzzi and Magoun, 1949). In recent studies, the selective optogenetic stimulation of some of these systems, including the noradrenergic neurons of the locus coeruleus and the dopaminergic neurons of the midbrain and dorsal raphe region, was sufficient to induce both EEG activation and behavioral arousal from sleep (Carter et al., 2010; Eban-Rothschild et al., 2016; Cho et al., 2017) or anesthesia (Taylor et al., 2016). Other recent studies have clarified the role of individual thalamic nuclei using electrical stimulation in monkeys (Redinbaugh et al., 2020; Bastos et al., 2021), and optogenetic stimulation in mice (Herrera et al., 2016; Honjoh et al., 2018). Arousal from NREM sleep and/or anesthesia occurs after stimulating thalamic nuclei with broad cortical projections, including the mouse ventromedial nucleus (VM) that projects to layer 1 of large parts of neocortex (Honjoh et al., 2018), as well as the monkey centrolateral nucleus that projects to superficial and deep layers of frontal and parietal cortex (Redinbaugh et al., 2020). By contrast, stimulation does not lead to arousal when directed at thalamic nuclei with more restricted projections, such as the mouse ventral posteromedial nucleus that connects to primary somatosensory cortex (Honjoh et al., 2018), the mouse ventral medial geniculate nucleus that projects to primary auditory cortex (Wang et al., 2023), and the monkey dorsomedial nucleus that is mainly connected to prefrontal cortex (Redinbaugh et al., 2020). Together, these results show that arousal from NREM sleep and/or anesthesia can be triggered from several distinct brainstem or thalamic nuclei, but only when their stimulation leads to broad activation of the cerebral cortex.

Whether arousal from unresponsive states can be obtained through direct activation of cortical neurons has not been tested. This is relevant given that the cerebral cortex is widely considered a central part of the neural substrate of consciousness and in most, although not all cases, consciousness is associated with responsiveness (Sanders et al., 2012). In fact, much of the current debate is not focused on whether the cortex contributes directly to consciousness but, rather, on whether this role can be ascribed to frontal or posterior cortical areas, or both (Boly et al., 2017; Odegaard et al., 2017). The reticular activating system and its components, on the other hand, are now generally viewed as supporting consciousness indirectly (Schiff, 2010; Koch et al., 2016), despite the fact that, in humans, lesions of the dorsolateral pontine tegmentum or paramedian midbrain usually result in immediate coma (Parvizi and Damasio, 2003; Posner and Plum, 2007). This is because patients with wide frontoparietal cortical network dysfunction typically remain unresponsive, in vegetative state, even when the function of the brainstem reticular formation is preserved (Laureys et al., 2004; Boly et al., 2008).

If the cortex is the core substrate of consciousness and the reticular activating system is only a “background condition” (Koch et al., 2016), it should be the case that the direct activation of cortical neurons is sufficient for arousal from unresponsive states, including from brainstem coma, when the function of the reticular activating system is impaired. Here we tested this hypothesis using CaMKIIα::ChR2 mice in which all cortical neurons can be optogenetically excited (see Methods), because our first goal was to assess in a direct and specific manner the overall role of the cerebral cortex. In the ongoing debate about which cortical regions contribute directly to consciousness, global workspace theory and higher-order theories ascribe a key role to fronto-parietal cortical networks (Dehaene and Changeux, 2011; Lau and Rosenthal, 2011), while others stress the role of posterior cortex (Lamme, 2006; Tononi et al., 2016). This issue remains highly debated (Boly et al., 2017; Odegaard et al., 2017). Here we selected the mouse medial prefrontal cortex (mPFC) and posterior parietal association cortex (PtA) or) to represent the “front” and “back” of the cortex, respectively. Although there are obvious limitations to the extent to which specific parts of mouse cortex can model higher order functions of the human cortex (Laubach et al., 2018), mPFC is essential for sensory processing, attention, planning, monitoring and emotion regulation (Le Merre et al., 2021) and PtA is a multisensory associative area implicated in perception, navigation and cognition (Whitlock, 2017; Lyamzin and Benucci, 2019).

Optogenetic stimulation of cortical neurons was performed during sleep, after induction of a coma-like state of unresponsiveness induced by the injection of the GABA_A_ receptor agonist muscimol in the midbrain reticular core, and during sevoflurane-dexmedetomidine (sevo-dex) anesthesia. Muscimol injections are likely to disfacilitate cortex and thalamus by removing the ascending arousal influence coming from the rostral reticular core (Minert and Devor, 2016; Lanir-Azaria et al., 2018). By contrast, sevo-dex anesthesia broadly affects brainstem, thalamus, hypothalamus and cortex both indirectly, mainly via disfacilitation caused by the block of noradrenaline release (dexmedetomidine), and directly, through GABA_A_-mediated inhibition (sevoflurane) (Hunter et al., 1997; Vinje et al., 2002; Sanders and Maze, 2012; Zhang et al., 2015; Iqbal et al., 2019).

We find that stimulation of either PtA or mPFC can quickly wake up mice from sleep and, when stronger and/or longer light pulses are used, can reverse the muscimol-induced, coma-like state, leading to both fronto-parietal EEG activation and recovery of the righting reflex (RORR). When the same mice are stimulated under deep sevo-dex anesthesia, EEG activation occurs in both cases without RORR. Thus, cortical activation and full arousal from unresponsive states such as sleep and “brainstem coma” can be triggered by direct stimulation of cortical neurons.

## Results

### Experimental design

Adult CaMKIIα::ChR2 mice of both sexes (> P56, n = 12, 5 females) were implanted with optic fibers for optogenetic stimulation of PtA or mPFC, intracortical laminar probes and surface electrodes for electroencephalographic (EEG) recordings, and cannulas aimed at the midbrain for muscimol injection (Fig. 1). Baseline 24-hour recordings of sleep and waking started at least a week after surgery, followed by stimulation experiments in which light pulses of different intensity were first administered during sleep and, later, during muscimol-induced coma and sevo-dex anesthesia.

**Figure 1.**
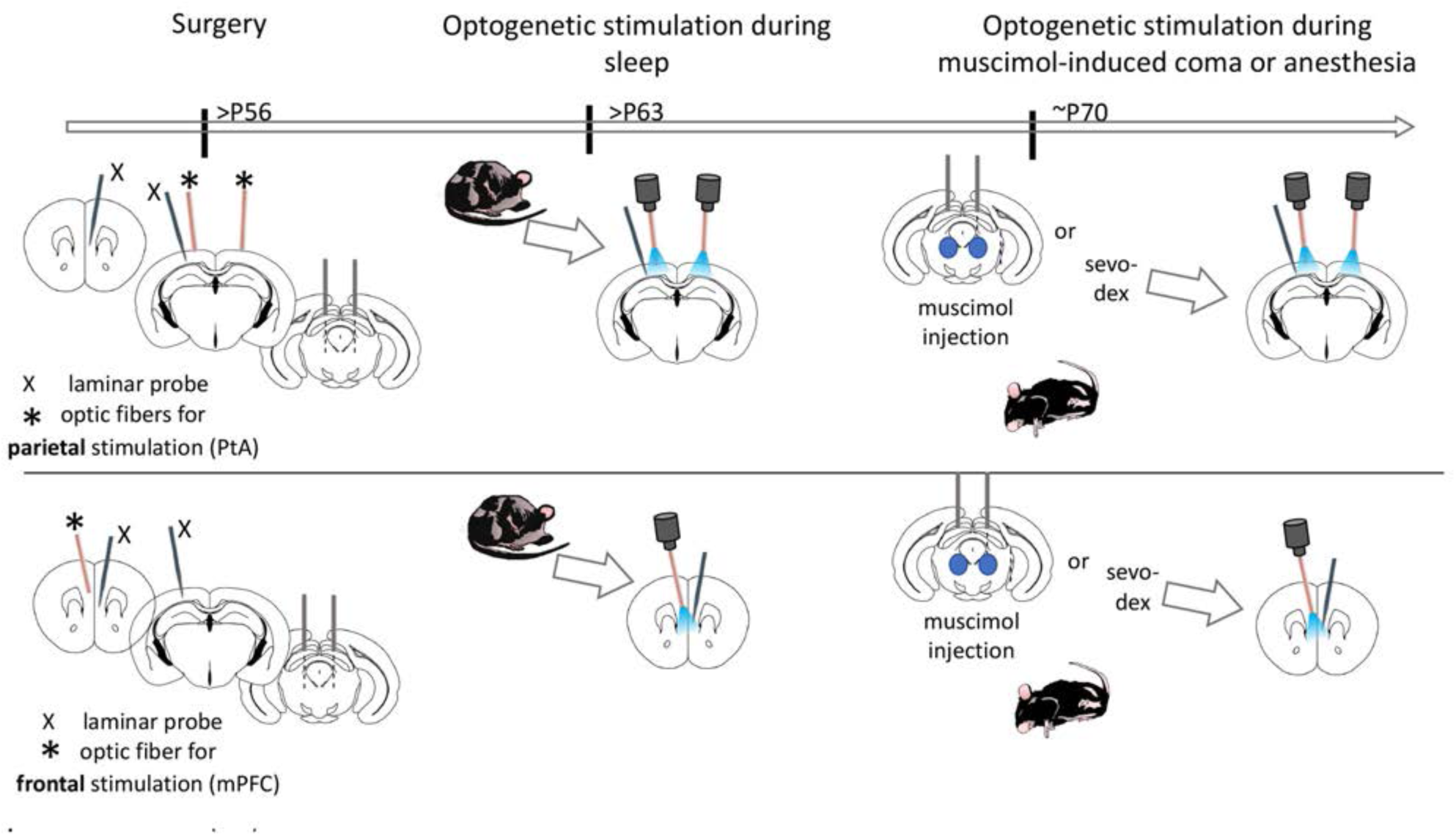
Experimental design and timeline. Mice (n = 12, 5 females) were implanted with two cannulas to deliver muscimol to the midbrain, 1-2 optic fibers above parietal association cortex (PtA, top row), 1 optic fiber close to the midline for bilateral optogenetic stimulation of medial prefrontal frontal (mPFC, bottom row; 4 mice) and laminar probes spanning frontal and parietal cortex (X). EEG electrodes (not shown) were also implanted. Two of the 8 mice with parietal stimulation had only one optic fiber (not shown). Optogenetic experiments occurred first during sleep and later after the induction of a coma-like state via muscimol injection in the midbrain or under sevo-dex anesthesia. Surgery and stimulation experiments were spaced at least one week apart.

CaMKIIα::ChR2 mice were obtained by crossing CaMKIIα-Cre mice with the Cre-dependent Ai32 strain, which expresses an improved channelrhodopsin-2/EYFP (ChR2-EYFP) fusion protein following exposure to Cre recombinase. Because the CaMKIIα promoter is broadly expressed in cortical glutamatergic neurons across areas and layers, the stimulation was expected to broadly excite the target area in both mPFC and PtA. Consistent with this, CaMKIIα::ChR2 mice showed broad cortical expression of ChR2-EYFP (Fig. 2A). For optogenetic experiments mice were implanted either with a single paramedian optic fiber inserted deep in the cortex to simultaneously target mPFC of both sides (Fig. 2B), or with two optic fibers over left and right PtA (Fig. 2C). A few mice had two fibers in left and right mPFC, or one fiber over left PtA (see below).

**Figure 2.**
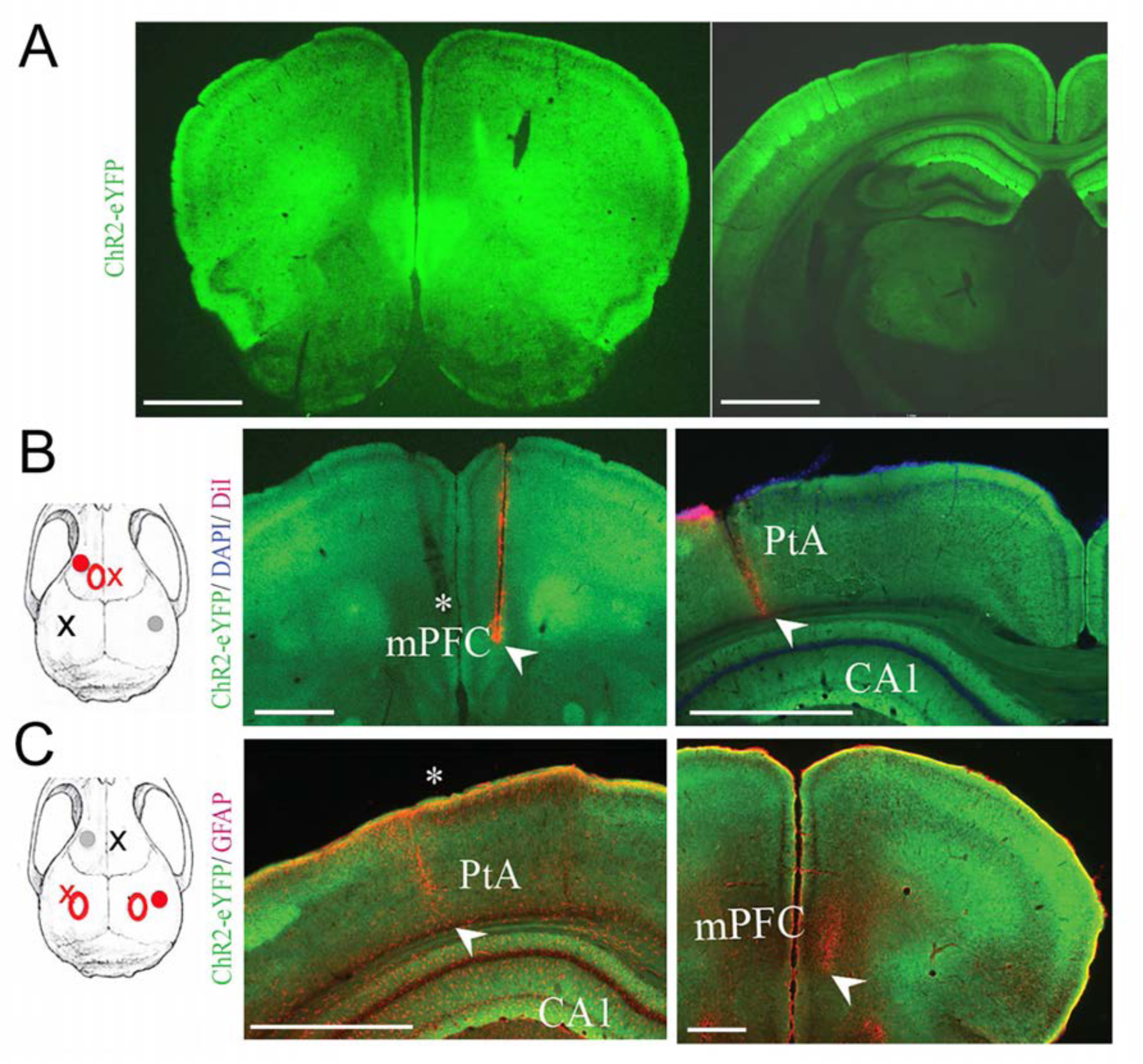
Immunohistochemical and histological analysis. **A,** representative coronal sections from one CaMKIIα::ChR2 mouse, confirming the broad cortical expression of ChR2-EYFP in frontal and parietal cortex. **B,** representative mouse with optogenetic stimulation in medial prefrontal cortex (mPFC). Left, dorsal schematic view of the mouse skull displaying the position of the optic fiber (red circle), laminar probes (X), and EEG screws (filled circles). Electrodes close to the optic fibers are in red. Middle and right, coronal sections showing the location of the tip of the optic fiber close to the midline in mPFC (*) and laminar probes in mPFC and posterior parietal cortex (PtA). The arrowheads indicate the tip of the laminar probes. DAPI and CM-DiI dye staining were used to identify cortical layers and probes, respectively. **C,** same as in B but for a representative mouse with optogenetic stimulation in PtA. In this case there were two optic fibers positioned above the cortical surface (one is shown, indicated by *). Coronal sections were stained with anti-GFAP (glial fibrillary acidic protein) antibody to identify the probes. Bars = 1 mm.

### Cortical optogenetic stimulation during sleep

In each CaMKIIα::ChR2 mouse laser pulses were delivered during NREM sleep using square pulses or 4-8 Hz train pulses lasting 1-5 sec. The different stimulation frequencies were meant to cover the broad range of firing rates typical of cortical pyramidal neurons (e.g. (Vyazovskiy et al., 2009; Hengen et al., 2016; Watson et al., 2016)) and our unpublished chronic Neuropixels recordings. The 0.1-0.5 Hz stimulation was a square pulse (e.g. 0.5 Hz, 1-sec per pulse) that elicits an initial peak photocurrent followed by a sustained plateau with decreased amplitude (Nagel et al., 2005; Lin et al., 2009; Berndt et al., 2011); by driving neurons to fire but not necessarily all at the same time, the pulse may promote a more physiological firing pattern.

In many cases, laser pulses were also delivered during rapid eye movement (REM) sleep. Stimulation experiments occurred over several days, and each day only a limited number of pulses was delivered, usually spaced minutes apart. During NREM sleep the arousal threshold varies depending on the amount of slow wave activity (SWA), which peaks at sleep onset and declines in the course of sleep (Neckelmann and Ursin, 1993). To control for the possible confound due to these homeostatic changes, stimulation experiments during sleep (and later during muscimol-induced coma or anesthesia) were performed approximately 5 to 8 hours after the beginning of the light phase, when most sleep pressure in mice has been released (Cavelli et al., 2023).

We applied cortical optogenetic stimulation during NREM sleep, when the EEG is dominated by slow waves that reflect the synchronous ON/OFF firing of cortical neurons (Steriade et al., 2001), and during REM sleep, when EEG pattern and cortical firing are similar to those of waking (Fig. 3A). Independent of the specific pattern or site of the stimulation (PtA or mPFC), mice always woke up from both NREM sleep and REM sleep (Fig. 3B,C). In every mouse, the awakening from REM sleep required significantly more laser power and/or longer stimulation compared to awakening from NREM sleep (p = 2.6e-5; Fig. 3C, Table 1). Moreover, across mice and independent of the pattern of stimulation, PtA stimulation was significantly more effective than mPFC stimulation, i.e. with PtA stimulation weaker and/or shorter pulses were needed to induce arousal (p = 7.7e-5; Fig. 3C, Table 1).

**Figure 3.**
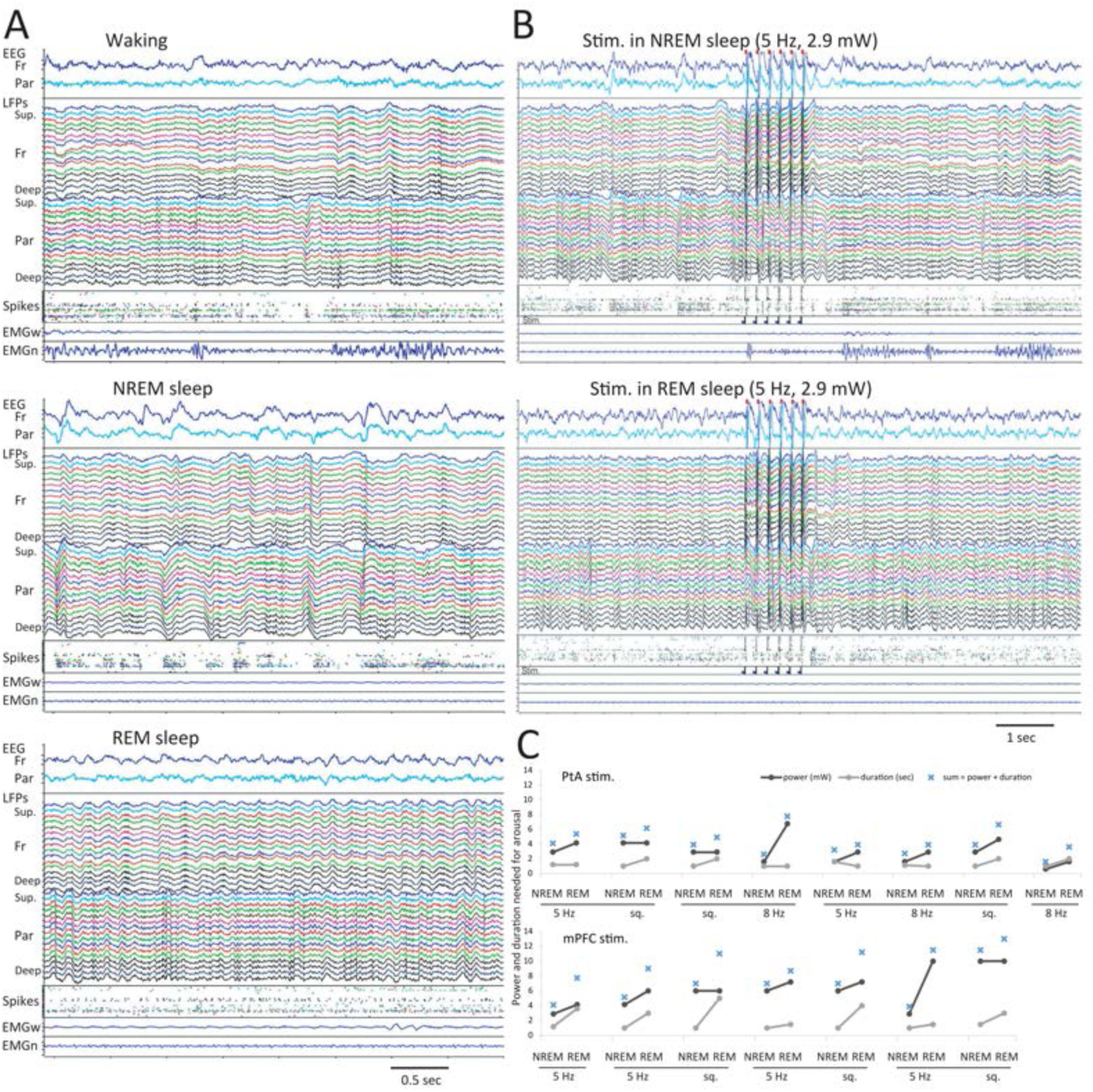
Optogenetic stimulation in sleep. **A,** representative traces (∼ 4 secs) of waking, NREM sleep, REM sleep in one representative mouse. For each behavioral state, the panel shows (from top to bottom) the electroencephalogram (EEG) from frontal and parietal cortex; local field potentials (LFPs) from prefrontal (Fr) and posterior parietal cortex (Par) recorded across layers with a laminar silicon probe (superficial on top) and thresholded spikes from the same LFP channels; electromyogram (EMG) from vibrissal (top) and neck musculature (bottom). LFPs and spikes from the same channel are color matched. **B,** example of the same optogenetic stimulation (prefrontal, bilateral stimulation; 5 Hz, 2.9 mW) leading to arousal from NREM sleep but not from REM sleep. Same mouse as in A. Stim. indicates when light pulses were given. **C,** Summary of the results of optogenetic stimulation during NREM sleep and REM sleep in each mouse. Horizontal bars below the x axis link data from the same mouse. In each experiment (NREM or REM sleep), the black and grey circle indicate, respectively, the minimum laser power (in mW) and the minimum length of stimulation (in sec) needed to wake up the mouse, based on 2-10 trials. Mice with optogenetic stimulation of posterior parietal association cortex (PtA) are shown in the top row, and mice with stimulation of medial prefrontal cortex (mPFC) are shown on the bottom row. In each experiment, the blue cross represents the sum of laser power (in mW) and duration (in sec). All 8 mice (4 for PtA, 4 for mPFC) received bilateral stimulation, either prefrontal (using one fiber implanted close to the midline) or posterior parietal (two fibers). Stimulation frequencies included 1-sec square pulses at 0.5Hz or trains (10ms; 5 or 8Hz).

**Table 1.** Summary of the results for the statistical tests presented in the manuscript, and sample size for figures 3, 5. Due to the unbalanced design with repeated measurements (Sample column), analyses are based on linear mixed effect models (lme4 package in R), with main effects tested using an asymptotic chi-squared test. Confidence intervals are based on an asymptotic z-test, and use the standard errors estimated from the LME model. The ratio of mean to variance is used as a measure of effect size, with the combined residual variance and random effect variance in the denominator (analogous to Cohen’s D but for LME models). Obs, observations (i.e. individual experiments).

| Result | Sample | p-value | 95% CI | Effect size |
| --- | --- | --- | --- | --- |
| State (REM, NREM) on needed stim. strength Fig. 3C | 8 mice, 28 obs | 2.6e-5 | (1.69, 3.90) | 1.88 |
| Position (PtA, mPFC) on needed stim. strength Fig. 3C | 8 mice, 28 obs | 7.7e-5 | (2.67, 5.26) | 2.66 |
| State (NREM, Coma) on Behav. score | 6 mice, 9 obs | 2.9e-13 | (-8.73, -7.27) | -7.98 |
| Position (PtA, mPFC) on LORR | 12 mice, 19 obs | 0.4844 | (-28.73, 13.53) | -0.42 |
| Position (PtA, mPFC) on pre-stim Behav. Score Fig. 5B and C | 12 mice, 19 obs | 0.8618 | (-1.29, 1.54) | 0.09 |

### Induction of a coma-like state in the mouse and behavioral analysis

In rats, the bilateral microinjection of GABA_A_ receptor agonists in a brainstem region called the mesopontine tegmental anesthesia area triggers an immediate and reversible state of profound unresponsiveness similar to coma or anesthesia (Devor and Zalkind, 2001). Although the exact mechanism underlying this state is complex, a likely candidate is a broad decline in excitability of the forebrain caused by the inhibition of the ascending reticular arousal system (Lanir-Azaria et al., 2018). In a first series of experiments, we tested whether we could induce a similar unresponsive state in mice. A total of 28 animals, including 14 CaMKIIα::ChR2 mice used in optogenetic experiments, 11 CaMKIIα-Cre mice, and 3 C57BL/6J mice, received a bilateral injection of the GABA_A_ receptor agonist muscimol in the mesencephalic reticular formation (Fig. 4A). All mice were briefly anesthetized with sevoflurane during the injections and the anesthesia was discontinued as soon as the procedure was completed. In a control experiment with saline injection, RORR occurred within 2 min from the time sevoflurane was discontinued, quickly followed by a normal cycling of sleep and wake episodes as during baseline. After muscimol injection, RORR also occurred within 1-2 min from the time the anesthetic was discontinued, but it was followed by an average period of around 40 minutes characterized first by hyperactivity with repetitive circling behavior, then by progressive ataxia followed by quiescence with the mouse lying on one side, and finally by loss of the righting reflex (LORR). In 5 mice in which no cortical optogenetic stimulation was performed after LORR, the period between LORR and spontaneous RORR was more than 2 hours (129 ± 12min; mean ± sem, 5 mice). During this period, the EEG pattern was dominated by large slow waves (Fig. 4B) and the mouse was breathing regularly, resting on the floor of the cage without any or with a few short spontaneous movements of the extremities.

**Figure 4.**
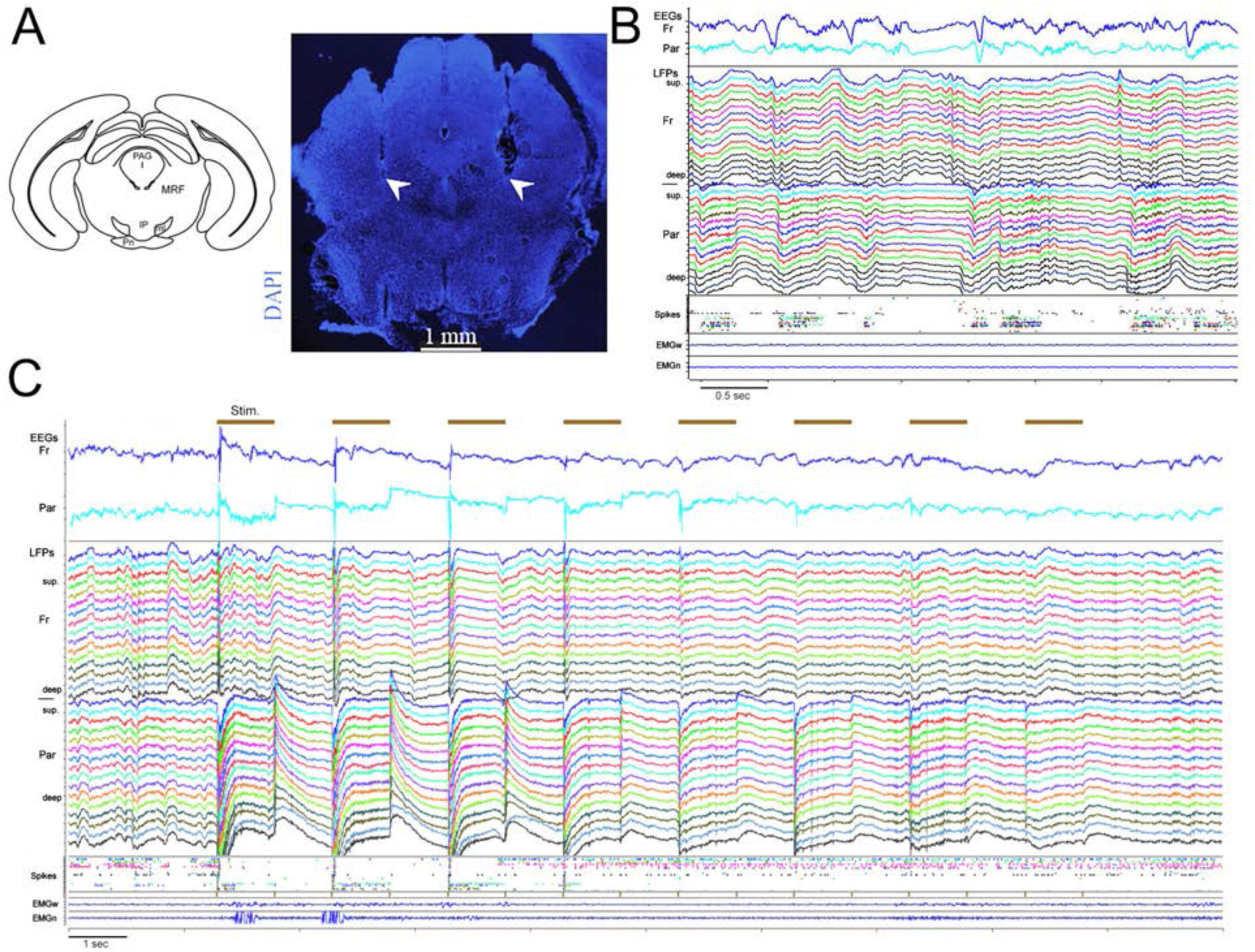
Muscimol-induced coma. **A,** schematic view of muscimol injection site and coronal section of a representative mouse showing part of the cannula tracks in the midbrain; arrowheads indicate the tip of the cannulas. IP, interpeduncular nucleus; ml, medial lemniscus; MRF, mesencephalic reticular formation; PAG, periaqueductal gray; pn, pontine nuclei. **B**, representative traces (∼ 4 secs) of cortical activity during muscimol-induced coma (after LORR). Same mouse as in Figure 3. The panel shows (from top to bottom) the electroencephalogram (EEG) from frontal and parietal cortex; local field potentials (LFPs) from frontal and parietal cortex recorded across layers with a laminar silicon probe (superficial on top) and thresholded spikes from the same LFP channels; electromyogram (EMG) from vibrissal (top) and neck musculature (bottom). LFPs and spikes from the same channel are color matched. **C**, example of the effect of PtA stimulation (8 pulses at 0.5 Hz, 2.9 mW) on EEG, LFP, and spike activity. Note the presence of slow waves in frontal and parietal LFPs, associated with bistable (on/off) firing (spikes) before stimulation, and EEG activation with tonic firing after the stimulation.

To establish the “depth” of this muscimol-induced state, we designed a battery of 6 stimuli that were delivered mostly in a fixed order, from mild to strong, before LORR (during NREM sleep), in the period between LORR and RORR (every 30-60 min), and after RORR. Previous behavioral scales in rats used only the standard righting reflex (Pal et al., 2018), and/or a series of behavioral tests to measure the responsiveness in more detail (Devor and Zalkind, 2001; Pais-Roldan et al., 2019). Our scale is in line with the ones developed for rats (Devor and Zalkind, 2001; Pais-Roldan et al., 2019), but expanded and optimized to be performed in freely moving mice. It also includes an olfactory test, which is informative given that olfaction is a predominant sensory modality in mice.

Based on the response to each stimulus we computed a cumulative score of responsiveness that could range from 0 (no response to any stimulus) to 12 (clear positive response to all 6 stimuli). When the stimuli were delivered during NREM sleep the total score was 12, that is, each stimulus triggered a positive response (score = 2) resulting in EEG activation and behavioral arousal (Fig 5A, 5B left). After muscimol-induced LORR instead, mice showed little (score = 1) or no response (score = 0) to most stimuli, although 5 mice still showed a positive response to odors (Fig. 5B and C). The cumulative score after muscimol-induced LORR was significantly reduced relative to NREM sleep (p = 2.9×10^-13^, Table 1). After spontaneous RORR (i.e. in experiments in which optogenetic stimulation did not occur), mice tried to stand on their paws and were drowsy, and the score remained below baseline levels for several hours, as shown in Fig. 5A for one representative mouse. By the next day, the responsiveness score, overall behavior and the sleep/wake pattern were back to normal. In experiments in which optogenetic stimulation occurred, the average cumulative score during the period between LORR and the first light pulse was 3.8 ± 1.5 (mean ± SD). In many cases, the same muscimol-induced state of unresponsiveness could be induced in the same animal 2-3 times, with experiments spaced approximately 1 week apart (9 mice). Based on this behavioral analysis we conclude that muscimol induces a state of long-lasting unresponsiveness that is deeper than NREM sleep. For simplicity, we call this state muscimol-induced “coma”.

**Figure 5.**
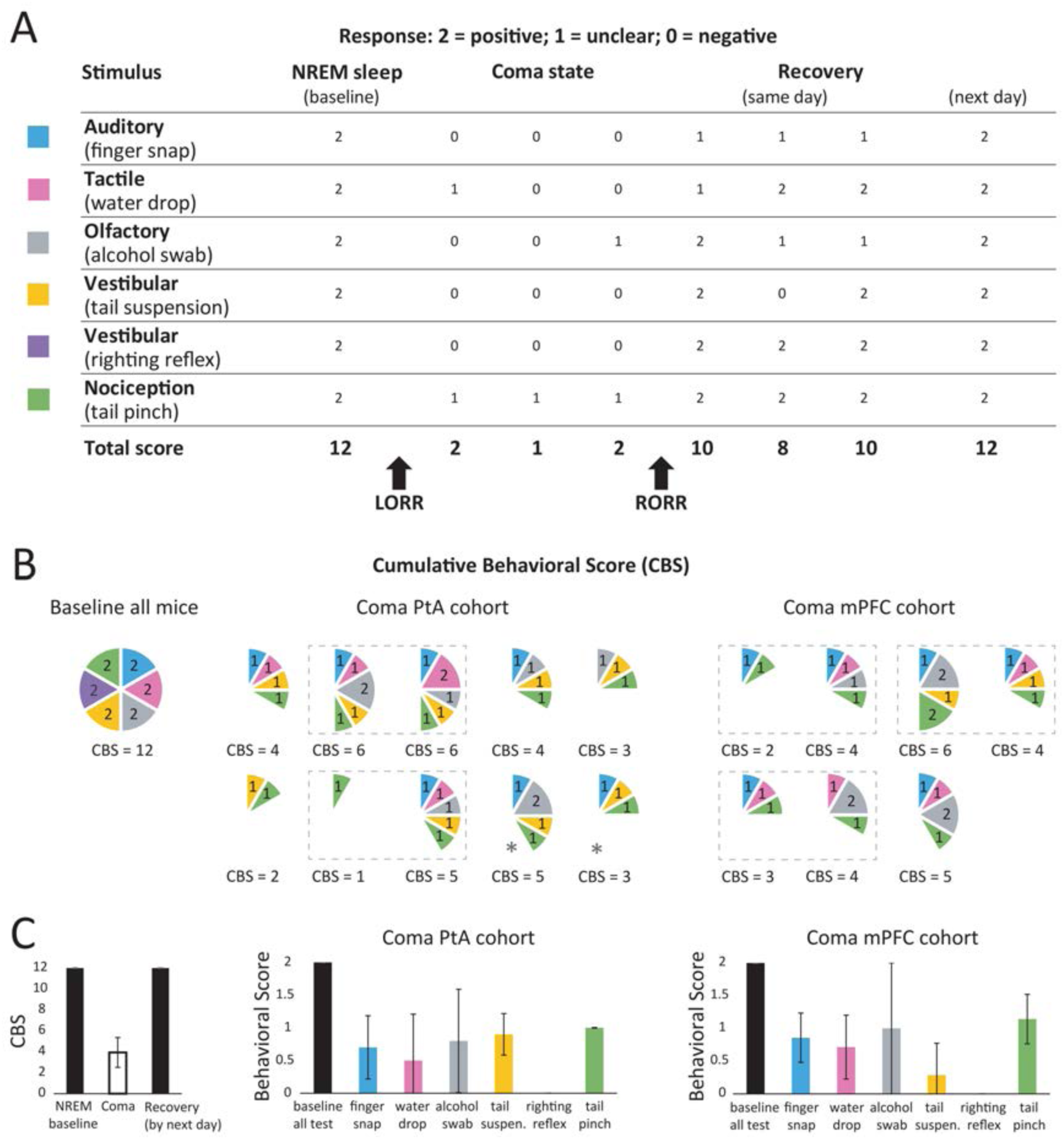
Behavioral characterization of muscimol-induced unresponsiveness. **A,** the behavioral test battery scores before, during and after the coma-like state in one mouse. LORR, loss of righting reflex. RORR, recovery of righting reflex. **B,** Pie charts showing the cumulative behavioral score (CBS) in baseline and during coma immediately before stimulation for all 12 mice used for optogenetic stimulation during coma. In the pie chart, the 6 stimuli are color-coded as shown in panel A. During baseline (left), all mice have the maximum CBS score of 12, i.e. they show a positive response (score = 2) to each of the 6 stimuli. During coma before stimulation, all mice fail to respond to the stimuli either partially (score = 1) or completely (score = 0), resulting in a decrease in CBS. Mice are grouped based on the site of the optogenetic stimulation (PtA, left, 8 mice; mPFC, right, 4 mice). Dashed frames link results of two coma experiments in the same mouse; * marks the mice with unilateral PtA stimulation. **C,** left, mean (± std dev) cumulative behavioral score (CBS) across all mice during NREM sleep, coma immediately before stimulation, and after recovery by the following day. Middle and right panels show mean (± std dev) scores across mice for each of the six stimuli.

### Cortical optogenetic stimulation during muscimol-induced coma

In 12 CaMKIIα::ChR2 mice the induction of a coma-like state was followed by cortical optogenetic stimulation in PtA or mPFC using square pulses (0.25 or 0.5Hz, lasting 1-2 sec) or trains (10ms; 4, 5, or 8Hz). Below, we describe these results separately for the two areas. Before the onset of the stimulation, the depth of the muscimol-induced coma was comparable in the two groups of animals. Specifically, latency to LORR (mean ± SD in min, PtA = 46.3 ± 21.0; mPFC = 41.0 ± 13.7; p = 0.48, Table 1) and behavioral score after LORR, immediately before the stimulation, did not differ between mPFC and PtA mice (total score, mean ± SD, PtA = 3.9 ± 1.7; mPFC = 4.0 ± 1.3; p = 0.86) (Fig. 5B and C, Table 1). In all mice the response to the vestibular stimulus (righting reflex) was negative.

Gamma power is often used as a proxy of neuronal firing (e.g. (Ray et al., 2008; Ray and Maunsell, 2011; Hayat et al., 2022)), and SWA power as an index of bistability (ON/OFF firing), which is associated with unconscious states (Casali et al., 2013). Thus, in each mouse, we measured high gamma power (70-100 Hz) during and after the light pulses to assess the immediate effects of the stimulation, and SWA (0.5-4 Hz) after the stimulation to test whether EEG activation had occurred (Fig. 4C). The mouse behavior was scored before, during, and after the stimulation in 4 categories (no/little movements, some movements, attempt to RORR, RORR).

#### PtA stimulation

In two mice carrying a single optic fiber over the left PtA light pulses (1 sec) were delivered every 2-4 seconds for a total of 10-20 sec. Optogenetic stimulation triggered immediately several movements of legs and body followed by attempts to right up (aRORR) after 30 or 60 sec from the onset of the stimulation, and full RORR in one of the two mice after 77 sec (Fig. 6A). Changes in gamma and SWA power were almost identical in the two mice. During the stimulation period, gamma power in the left parietal LFP electrode, the closest to the stimulated site, increased when the light was on, with little change in the other electrodes (Fig. 6A). EEG activation was evident on the stimulated side, with a large decrease in SWA in the left parietal LFP electrode and a smaller decrease in the left frontal electrode, while there were no changes contralaterally (Fig. 6A).

**Figure 6.**
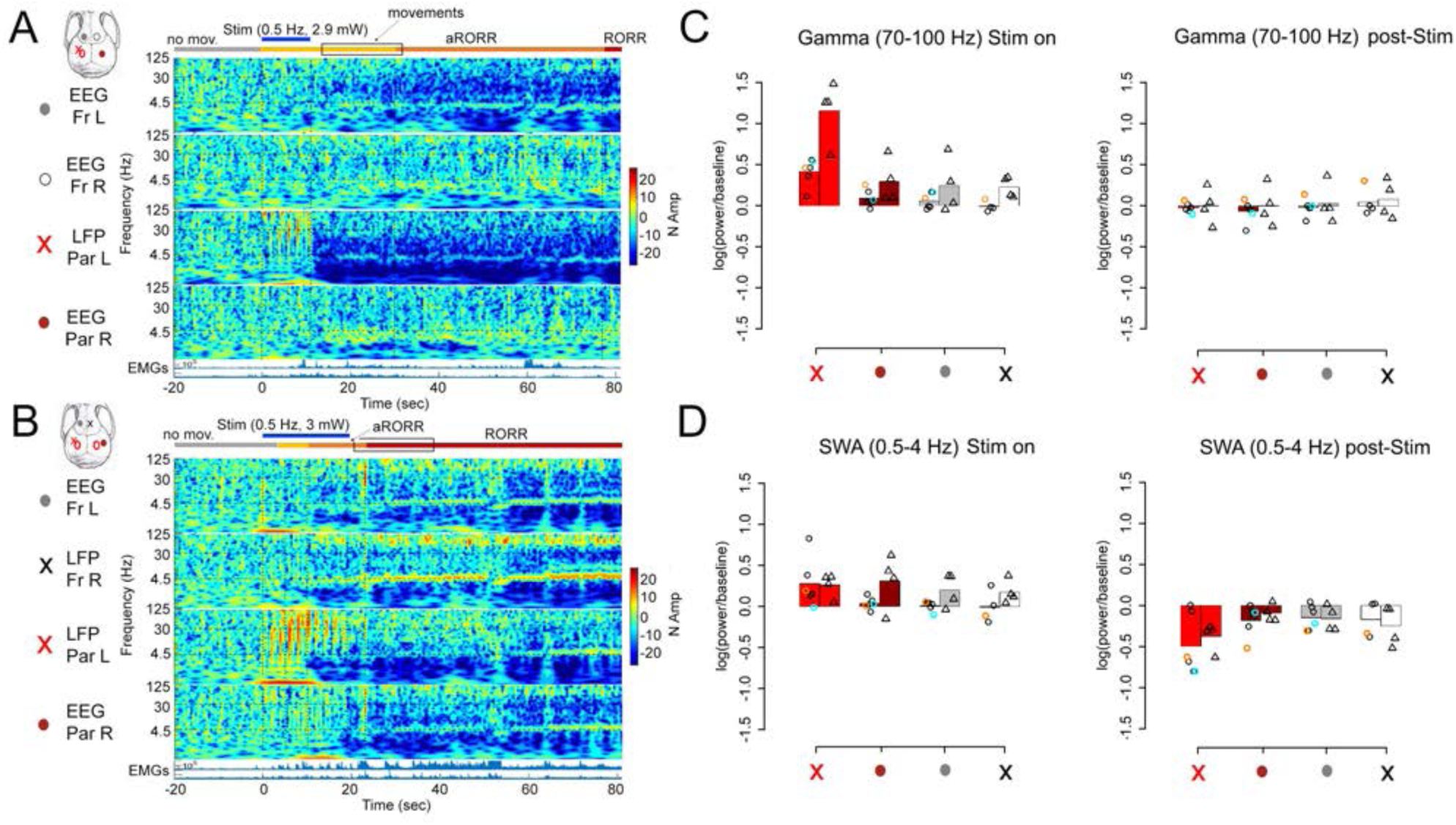
Optogenetic stimulation of PtA during muscimol-induced coma. **A,** mouse implanted with a single optic fiber (red circle) over left PtA just medial to the laminar probe (red X) (unilateral parietal stimulation). Relative power spectrum for each of the 4 electrodes (3 EEGs indicated by circles, 1 deep LFP channel indicated by X) starting 20 sec before the stimulation (time 0), when the mouse was in a stable coma-like state. The power for each frequency bin (0.0078Hz) is normalized based on the last 50 to 1 sec before the first laser pulse. EMGs, electromyograms from vibrissal (top) and neck musculature (bottom). The duration of the stimulation is indicated by the blue horizontal bar on top, followed by a rectangle indicating the post-stimulation time window when SWA was measured. Behavior was scored using videorecording and is indicated by horizontal bars; grey bars indicate when no/few movements were present; yellow: more movements of limbs and body; orange: attempts to RORR (aRORR); red: RORR. **B,** same as in A for a mouse implanted with two optic fibers (red circles) over left and right PtA (bilateral parietal stimulation). In this mouse the stimulation at 0.5 Hz triggered EEG activation and RORR. **C** and **D**, bar plots indicating the mean change in gamma power and SWA during and after stimulation, with individual experiments indicated. Left columns with circles indicate low frequency stimulation, and right columns with triangles indicate high frequency stimulation. The experiments shown in panels A and B are highlighted in cyan and orange, respectively.

In six mice carrying two optic fibers over left and right PtA, optogenetic stimulation led to aRORR within 5-25 secs from the onset of stimulation, followed by RORR (latency from stimulation onset 23-174 sec). Consistent with the findings with unilateral PtA stimulation at 0.5 Hz, gamma power increased during the stimulation in the parietal electrodes, especially the left parietal LFP close to the optic fiber (Fig. 6B). EEG activation (SWA decrease) was prominent post-stimulation in the parietal electrodes and also clearly present in the frontal electrodes (Fig. 6B). While the stimulation at 0.5 Hz was effective in most cases, it failed to awake two animals. In both mice, the stimulation was maintained for more than 1 minute, but it only triggered some movements that never evolved into aRORR or RORR. In these two cases the increase in gamma power was small in the parietal electrodes, and SWA did not change. However, aRORR/RORR could be triggered in both mice using pulses at 4 or 5 Hz rather than at 0.5 Hz. Across all experiments with PtA stimulation (n = 10 experiments in 8 mice), aRORR/RORR occurred in 67% of the cases (4/6 experiments) with pulses at 0.5 Hz or lower (“low frequency stimulation”) and in 100% of the cases with pulses at 4 or 5 Hz (“high frequency stimulation”). Figure 6C,D shows the changes in gamma power and SWA during and after the stimulation for each individual experiment, as well as mean changes: in general, changes seemed most prominent in the left parietal LFP (closest to the optic fiber).

#### mPFC stimulation

In the first two mice two optic fibers were implanted, one in the mPFC of each side, and in both animals optogenetic stimulation at 5 Hz quickly induced RORR from muscimol-induced coma, associated with a large increase in gamma power mainly in the frontal LFP electrode and clear signs of EEG activation. However, the histology revealed that in both animals the tip of the fibers was too deep and reached the white matter and therefore these mice were excluded for all analyses. To avoid this problem, in the next 4 mice a single fiber was implanted in the left mPFC more rostrally and close to the midline, to allow the light pulses to also reach the contralateral side (Fig. 2B). In all four mice histology confirmed the position of the optic fiber within the prefrontal grey matter. In two of these animals the stimulation at 0.5 Hz failed to induce signs of EEG activation and RORR (Fig. 7A). Stimulation at 5 Hz instead induced aRORR/RORR in 80% of cases (4/5 experiments), confirming that stimulation at 5 Hz was more effective than at 0.5 Hz. In 2 of the 4 mice abnormal, hypersynchronous activity occurred for several seconds after the stimulation and then subsided (Fig. 7B). Across all experiments with mPFC stimulation (n = 7 experiments in 4 mice), aRORR/RORR and/or EEG activation never occurred with pulses at 0.5 Hz, while they did happen in 80% of the cases with pulses at 5 Hz. Figure 7C,D shows the changes in gamma power and SWA during and/or after the stimulation for each individual experiment, as well as mean changes: during the stimulation changes in gamma power seemed most prominent in the local LFP channel, while no obvious effects on SWA were present after stimulation.

**Figure 7.**
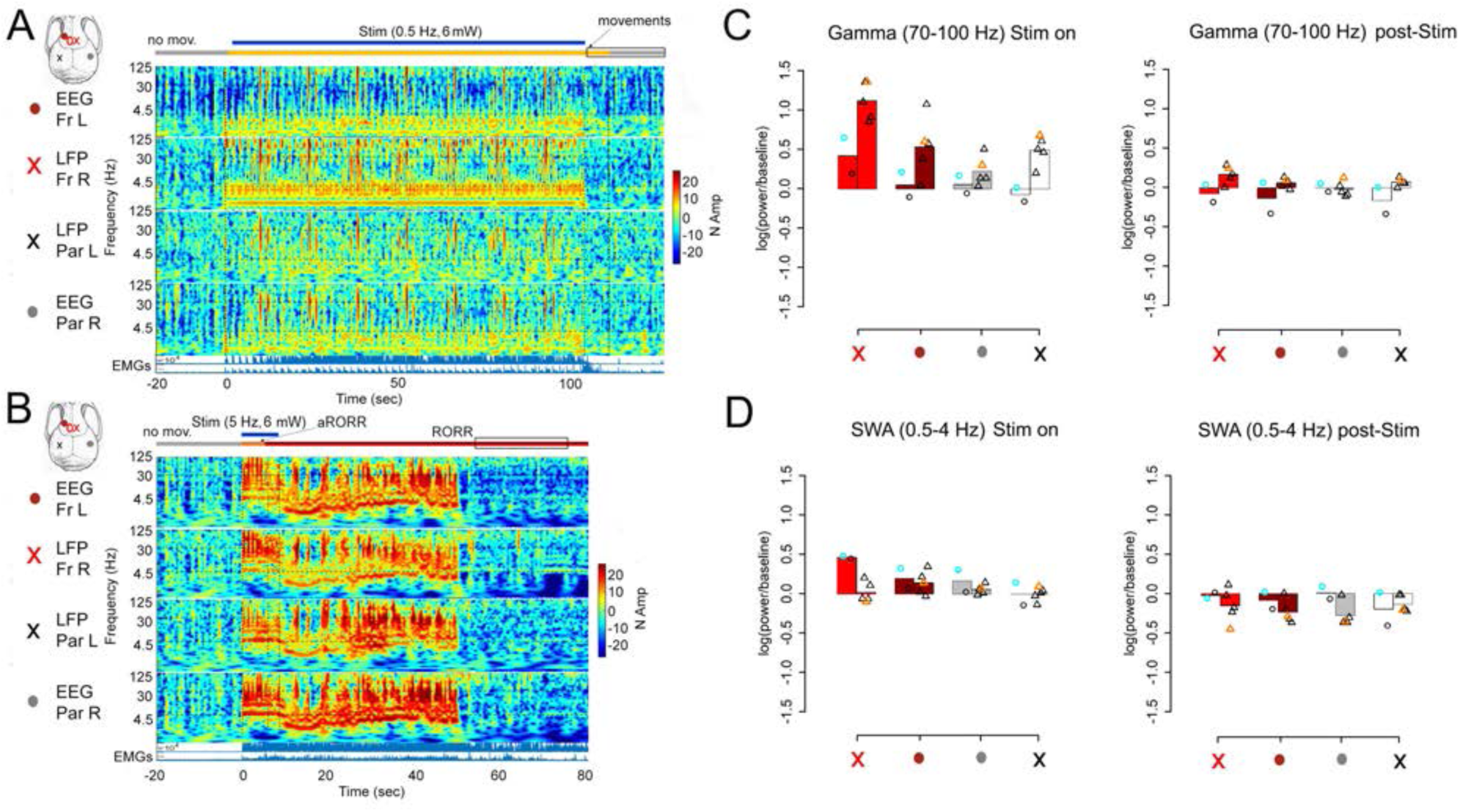
Optogenetic stimulation of mPFC during muscimol-induced coma. **A,** a mouse implanted with one optic fiber close to the midline (red circle) for bilateral prefrontal stimulation. In this mouse the stimulation at 0.5 Hz triggered neither RORR nor EEG activation. Relative power spectrum for each of the 4 electrodes (2 EEGs indicated by filled circles, 2 deep LFPs indicated by X). Time 0 indicates stimulation onset, which occurs when the mouse was in a stable coma-like state. The power for each frequency bin (0.0078Hz) is normalized based on the last 50 to 1 sec before the first laser pulse. EMGs, electromyograms from vibrissal (top) and neck musculature (bottom). The duration of the stimulation is indicated by the blue horizontal bar on top, followed by a rectangle indicating the post-stimulation time window when SWA was measured. Behavior was scored using videorecording and is indicated by horizontal bars; grey bars indicate when no/few movements were present; yellow: more movements of limbs and body; orange: attempts to RORR (aRORR); red: RORR. **B**, same mouse as in A. The stimulation at 5 Hz triggered RORR and EEG activation, but hypersynchronous activity was obvious in all electrodes after the stimulation ended. SWA was measured at the end of this abnormal activity (∼ 40 sec after the end of stimulation). **C** and **D**, bar plots indicating the mean change in gamma power and SWA during and after stimulation, with individual experiments indicated. Left columns with circles indicate low frequency stimulation, and right columns with triangles indicate high frequency stimulation. The experiments shown in panels A and B are highlighted in cyan and orange, respectively.

### Stimulation during anesthesia

Optogenetic stimulation was performed during deep sevo-dex anesthesia (1-2% sevo, 70-100ug/kg dex, n= 9 mice), while cortical activity was dominated by highly synchronous, large slow waves (Fig. 8A). The combination of sevo-dex was chosen to maintain a stable level of slow-wave anesthesia, long enough to allow for the optogenetic stimulation. Sevo has low blood solubility and fast pharmacodynamics and in our experience, when given alone, is either unable to generate a steady level of anesthesia (at low dose) or leads to burst-suppression (at high dose).

**Figure 8.**
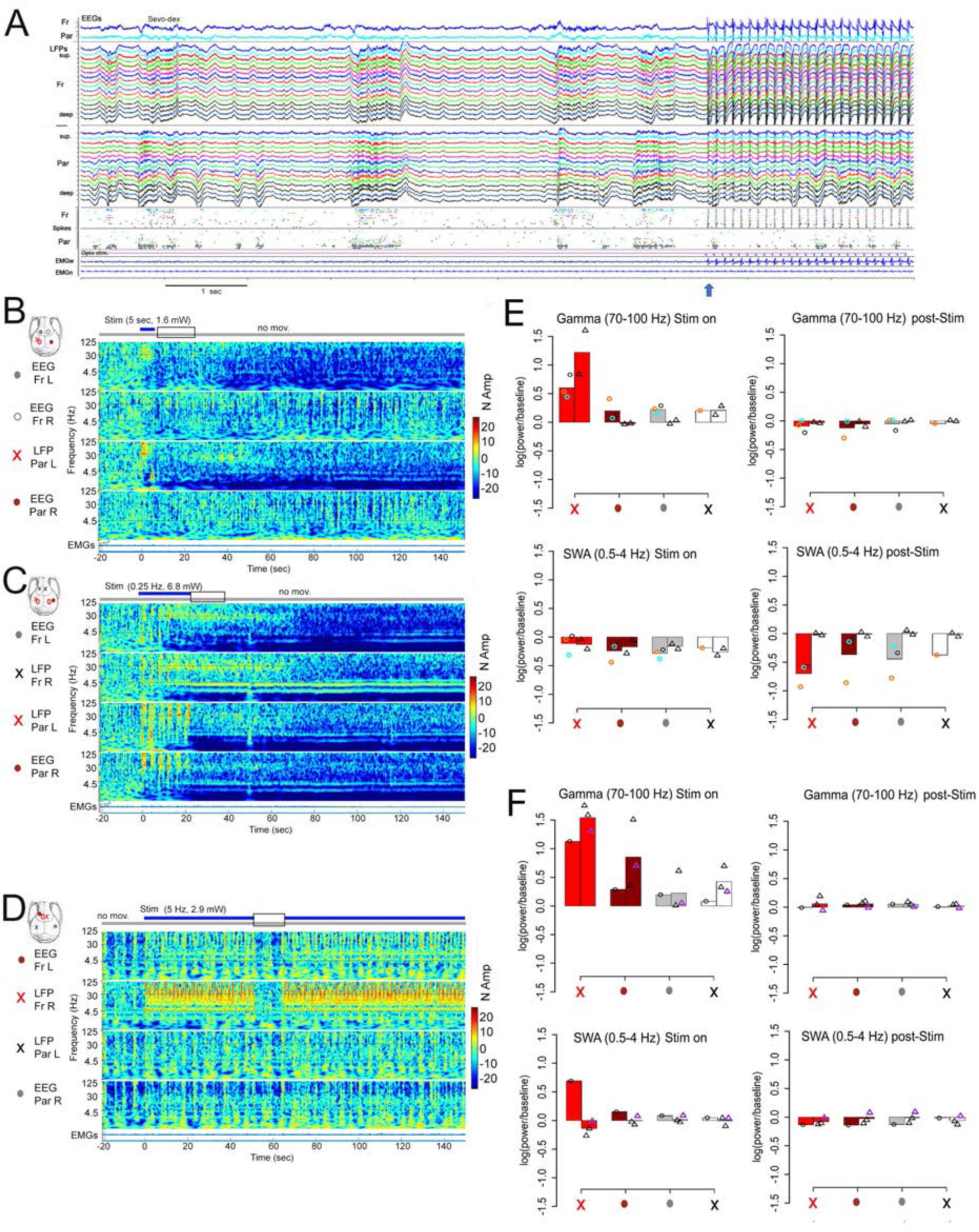
Optogenetic stimulation during sevo-dex anesthesia. **A**, representative traces of cortical activity during stable sevo-dex anesthesia (after LORR), one minute before stimulation (left) and just before and during stimulation (right). Blue arrow indicates the onset of the light pulses (25 pulses are shown). Same mouse as in Figure 3A (prefrontal, bilateral stimulation). The panel shows (from top to bottom) the electroencephalogram (EEG) from frontal and parietal cortex; local field potentials (LFPs) from frontal and parietal cortex recorded across layers with a laminar silicon probe (superficial on top) and thresholded spikes from the same LFP channels; electromyogram (EMG) from vibrissal (top) and neck musculature (bottom). **B,** unilateral stimulation in PtA. Left, relative power spectrum for each of the 4 electrodes (3 EEGs indicated by circles, 1 deep LFP indicated by X). The electrode close to the optic fiber is shown in red. Time 0 indicates stimulation onset. The power for each frequency bin (0.0078Hz) is normalized based on the last 50 to 1 sec before the first laser pulse. EMGs, electromyograms from vibrissal (top) and neck musculature (bottom). The duration of the stimulation is indicated by the blue horizontal bar on top, followed by a rectangle indicating the post-stimulation time window when SWA was measured. Behavior was scored using videorecording and is indicated by horizontal bars (grey bars indicate when no/few movements were present). **C**, as in B for a mouse with bilateral stimulation in PtA. Left, relative power spectrum for each of the 4 electrodes (2 EEGs indicated by filled circles, 2 deep LFPs indicated by X). **D,** as in B and C in a mouse with stimulation in mPFC. **E**, bar plots indicating the mean change in gamma power and SWA during and after PtA stimulation, with individual experiments. Left columns with circles indicate low frequency stimulation, and right columns with triangles indicate high frequency stimulation. The experiments shown in panels B and C are highlighted in cyan and orange, respectively. **F**, as in E, after mPFC stimulation. The experiment shown in panel D is highlighted in purple.

Mice were lying on their side, and the total score on the behavioral battery performance was the lowest possible (total score = 0) and identical for mice that received PtA or mPFC stimulation. Stimulation in either PtA or mPFC never resulted in aRORR or RORR even when light pulses delivered during sevo-dex anesthesia were stronger and/or longer than those applied during muscimol-induced coma. In the two mice carrying a single optic fiber, unilateral PtA stimulation resulted in clear EEG activation (SWA decrease) in both frontal and parietal cortex of the stimulated side (Fig. 8B), while bilateral strong PtA stimulation resulted in a broad increase in gamma power and EEG activation in all channels (Fig. 8C). In all PtA experiments (5 experiments in 4 mice) an increase in gamma power during the stimulation, and a decrease in SWA after the stimulation, were apparent in the local LFP channel (Fig. 8E). In the 4 mice carrying one fiber targeting mPFC (4 experiments in 4 mice), an increase in gamma power was apparent in the frontal electrodes close to the optic fiber, with no obvious effects on SWA after the stimulation (Fig. 8D,F).

## Discussion

In this study we used optogenetic stimulation to assess the ability of cortical pyramidal neurons to trigger cortical activation and behavioral arousal from sleep, muscimol-induced coma, and sevo-dex anesthesia, three states of progressively deeper unresponsiveness as measured using a behavioral test battery.

Cortical optogenetic stimulation could always awaken the mice from NREM sleep. This was also the case for REM sleep, although arousal from this phase required stronger and/or longer light pulses compared to NREM sleep. A candidate mechanism that could partially account for these results is the level of activity of the noradrenergic system of the locus coeruleus (LC), because LC activity is important for arousability and the LC is actively inhibited during REM sleep. Specifically, optogenetic stimulation of the LC invariably awakens the mice with short latency from both NREM sleep and REM sleep (Carter et al., 2010). Relative to controls, mice unable to produce noradrenaline require more noise to wake up from recovery sleep following sleep deprivation (Hunsley and Palmiter, 2004) and in rats, optogenetic silencing of LC neurons reduces the likelihood of sound evoked awakenings (Hayat et al., 2020). LC noradrenergic neurons fire maximally during waking, much less so in NREM sleep and not at all during REM sleep (Aston-Jones and Bloom, 1981a; Takahashi et al., 2010; Hayat et al., 2020), when LC neurons are actively inhibited (Nitz and Siegel, 1997; Verret et al., 2006; Luppi et al., 2011). Recent studies in mice using genetically encoded noradrenaline sensors (Feng et al., 2019) also showed that during NREM sleep noradrenaline levels continue to fluctuate up and down every 30-50 seconds in both thalamus (Osorio-Forero et al., 2021) and prefrontal cortex (Kjaerby et al., 2022), while they steadily decline in these regions during REM sleep. Intriguingly, other optogenetic experiments that targeted the thalamus found even more extreme differences in arousability between NREM sleep and REM sleep. Bilateral optogenetic excitation of the matrix cells of the ventromedial thalamic nucleus (VM), which sends diffuse glutamatergic projections to layer 1 of neocortex, could wake up the mouse from NREM sleep but not from REM sleep (Honjoh et al., 2018). A similar result was observed after bilateral optogenetic inhibition of the GABAergic cells of the reticular thalamic nucleus, which strongly inhibits the rest of the thalamus (Takata, 2020). Of note, when arousal threshold is measured using peripheral (acoustic) stimuli, arousal from deep NREM sleep (slow wave sleep) requires louder stimuli than from REM sleep, in both humans and rodents (Rechtschaffen et al., 1966; Zepelin et al., 1984; Neckelmann and Ursin, 1993), although when tonic REM sleep and phasic REM sleep are tested separately, the latter is as deep as slow wave sleep (Ermis et al., 2010). Also, the scent of a predator wakes up a mouse more rapidly from REM sleep than from NREM sleep (Tseng et al., 2022). In response to a mild sound, LC unit activity strongly increases during wake and not at all in REM sleep, while during NREM sleep the evoked firing response is small but still present (Aston-Jones and Bloom, 1981b). Thus, in physiological conditions additional mechanisms must exist to regulate arousability from NREM sleep. Among them, the ON/OFF bistable pattern of activity in the thalamocortical system, which is responsible for the occurrence of slow waves, is a primary candidate because it disrupts thalamocortical and corticocortical connectivity (Massimini et al., 2005). In line with this, arousal thresholds within NREM sleep are positively correlated with SWA (Neckelmann and Ursin, 1993; Hayat et al., 2020).

The direct excitation of cortical cells could also revert the state of unresponsiveness caused by muscimol injection in the midbrain. Cortical activity switched back to a wake-like, tonic pattern of firing and mice could stand up and walk, even if the midbrain was directly inhibited. Patients do not recover consciousness if frontoparietal cortex is broadly damaged, even when the brainstem is intact (Laureys et al., 2004; Boly et al., 2008). Our results therefore complement clinical findings and provide independent evidence for the key role of the cortex, but not the brainstem, in supporting consciousness. During muscimol-induced coma, behavioral arousal as measured by RORR could be triggered by stimulation of either posterior parietal cortex or medial prefrontal cortex. In all successful cases the stimulation quickly recruited both frontal and parietal regions. This is consistent with the results of thalamic electrical stimulation in anesthetized monkeys, in which behavioral arousal occurred after activation of the centrolateral nucleus, which projects to both frontal and parietal cortex, but not after stimulation of the dorsomedial nucleus, which is mainly connected to prefrontal cortex (Redinbaugh et al., 2020).

During sevo-dex anesthesia the direct optogenetic stimulation of cortical cells produced cortical activation but did not result in behavioral arousal (RORR). Dexmedetomidine is a highly selective α2-adrenergic receptor agonist that causes sedation and, at higher doses, LORR (Zhang et al., 2015). It acts through several pre- and postsynaptic mechanisms, including the widespread block of noradrenaline release and the direct local inhibition of LC neurons (Aghajanian and VanderMaelen, 1982; Sanders and Maze, 2012; Zhang et al., 2015). Sevoflurane broadly inhibits neuronal activity mainly by acting as a positive modulator of the GABA_A_ receptor, although it also antagonizes excitatory NMDA receptors, promotes two-pore domain potassium conductances, and blocks glutamate release (Vinje et al., 2002; Iqbal et al., 2019). Together, these drugs have profound depressing effects on most of the brain—not only on the cerebral cortex, but also on thalamus, basal ganglia, and brainstem. Cortical activation in the absence of behavioral arousal is characteristic of REM sleep, a behavioral state of quiescence and reduced responsiveness almost always accompanied by dreaming. When humans dream, whether during REM sleep or NREM sleep, cortical activation, indexed like here by decreased slow wave activity, is observed primarily in posterior cortex, whereas prefrontal cortex remains deactivated (Siclari et al., 2017; Bernardi et al., 2019). Preserved consciousness accompanied by unresponsiveness or minimal responsiveness is also observed in neurological conditions, especially when associated with massive lesions of prefrontal-basal ganglia-thalamic circuits and of the dopaminergic system ((Schiff et al., 2014; Casarotto et al., 2016; Boly et al., 2017); for a striking example of akinetic mutism, see (Comanducci et al., 2023)). Thus, the induction of cortical arousal without behavioral arousal in our deep sevo-dex anesthesia condition may be due to the inability of cortical stimulation to activate prefrontal-basal ganglia-thalamic circuits, which seems to be one of the mechanisms that distinguish disconnected from connected consciousness (Sanders et al., 2012; Comanducci et al., 2023).

The lack of RORR with bilateral cortical stimulation under sevo-dex anesthesia contrasts with the results of the bilateral optogenetic stimulation of VM (Honjoh et al., 2018). In that study we found that all 6 mice anesthetized with sevo-dex showed cortical EEG activation, 5 of them exhibited continuous limb movements, and 4 had full RORR within 1-4 minutes from the onset of VM stimulation (Honjoh et al., 2018). The lower doses of sevoflurane and dexmedetomidine in that study (sevo: 1-1.2%; dex: 50-70 ug/kg) are unlikely to account for the different outcome of the stimulation because both studies used the minimum dose required for a stable slow wave anesthesia (in different ambient temperatures and mouse strains). Instead, a key factor may be that while here we stimulated a single cortical area, VM stimulation can broadly activate many cortical regions, as well as basal ganglia and mesencephalic circuits that are important for behavioral arousal. Anatomically, VM neurons are multi-area matrix cells that project to layer 1 of most of the neocortex and further innervate the central region of the caudate-putamen and midbrain tegmentum (Herkenham, 1979). Intriguingly, VM stimulation induces RORR from sevo-dex anesthesia but not from REM sleep (Honjoh et al., 2018). The only area spared by VM axons is the retrosplenial cortex (Herkenham, 1979), which is increasingly recognized as a main cortical hub for the generation and maintenance of REM sleep (Dong et al., 2022; Wang et al., 2022).

In contrast with our results, a recent study in rats anesthetized with sevoflurane (Pal et al., 2018) concluded that “consciousness” could be restored by pharmacological activation of prefrontal prelimbic cortex (our mouse mPFC) but not posterior and medial posterior cortex (our mouse PtA). This conclusion was based on the fact that, while the injection of carbachol or noradrenaline in any of these areas triggered EEG activation and increased respiratory rate, only the pharmacological activation of prefrontal prelimbic cortex resulted in RORR in 4 out of 11 rats (Pal et al., 2018). These results cannot be directly compared to ours due to differences in species (rats vs mice), anesthesia (sevo vs sevo-dex), method of cortical stimulation (pharmacological vs optogenetic), and length of the stimulation. The last factor is especially important, because the drugs were infused continuously for 12.5 minutes and aRORR/RORR were observed after several minutes, making it difficult to rule out subcortical effects due to the diffusion of the drug. Of note, optogenetic stimulation of the dopaminergic axons targeting mPFC does not induce arousal from NREM sleep, while stimulation of dopaminergic fibers targeting the nucleus accumbens and the dorsolateral striatum does (Eban-Rothschild et al., 2016). Independent of the possible role of subcortical areas, it was the presence or absence of RORR and wake-like motor behavior that prompted the authors to conclude that prefrontal, but not posterior, cortex plays a key role in restoring “signs of consciousness” (Pal et al., 2018). Equating consciousness with behavioral arousal, however, ignores the substantial evidence that consciousness can be present in unresponsive states during which posterior cortical regions are activated, as is the case with REM sleep and widespread prefrontal lesions. Indeed, in the Pal et al. study, pharmacological activation of posterior and medial posterior cortex triggered EEG activation with increased theta to SWA ratio, increased respiratory rate, increased cortical levels of acetylcholine, and in some cases muscle twitches. In the absence of large movements and locomotion, this combination of features is typical of REM sleep. Unlike humans, mice do not report whether they had been dreaming after awakenings from REM sleep. However, the perturbational complexity index--the most sensitive and specific maker of consciousness, validated in humans across many conditions of consciousness and unconsciousness—is similarly high in wakefulness and REM sleep in both humans (Massimini et al., 2010; Casarotto et al., 2016) and rodents (Cavelli et al., 2023). This further emphasizes the importance of distinguishing between consciousness and responsiveness (Sanders et al., 2012; Koch et al., 2016; Comanducci et al., 2023).

The present results show that awakening from both NREM and REM sleep through optogenetic stimulation required significantly weaker and/or shorter light pulses in parietal cortex than in prefrontal cortex. We also observed that PtA stimulation resulted in a greater proportion of RORR from muscimol-induced coma than mPFC stimulation. However, direct comparisons between PtA and mPFC experiments are not possible due to the unbalanced nature of the experiments, which used different frequencies and patterns of stimulations that were not matched between parietal and frontal stimulations. We also cannot prove that the strength of the optogenetic stimulation was perfectly matched in the two groups of mice.

Because animals were exclusively implanted with cortical electrodes, we could not assess the effects of optogenetic cortical stimulation on subcortical structures. Due to the duration of the stimulation paradigm (several seconds), cortical stimulation could have indirectly activated both diencephalic and brainstem nuclei. Thus, it is possible that cortical and behavioral arousal may have been mediated by the direct activation of cortico-cortical networks, the recruitment of cortico-thalamo-cortical loops, or the secondary involvement of brainstem circuits. However, the cortical and behavioral arousal from the deep coma induced by muscimol injections in the brainstem suggests that cortical stimulation awakened the mice despite the pharmacological suppression of brainstem circuits that normally control arousal. Thus, cortico-thalamic networks may be sufficient to autonomously support a conscious state, while brainstem arousal systems provide facilitating background conditions that can be substituted for by direct cortical activation.

This descriptive study has several limitations, starting from the small number of mice and the unbalanced design of the experiments. Also, we only targeted one site in the front and one in the back of the cortex, thus we do not know whether the conclusions would extend to other frontal and parietal areas. Finally, our optogenetic manipulation affects most, if not all, cortical neurons. An important next step would be to test the role of select populations of pyramidal neurons, as well as the role of specific cortical layers in supporting consciousness.

## STAR METHODS

### 1. RESOURCE AVAILABILITY

- **Lead contact.** Additional information and requests for resources and reagents should be sent and will be fulfilled by the lead contact, Chiara Cirelli.
- **Materials availability.** This study did not generate new unique reagents.
- **Data and code availability.** Data reported in this paper will be shared by the lead contact upon request. This paper does not report original code but the analysis scripts are available (https://github.com/maomrong/mao-2023). Any additional information required to reanalyze the data reported in this paper is available from the lead contact upon request.

### 2. EXPERIMENTAL MODEL AND SUBJECT DETAILS

#### Experimental animals

Adult mice (CaMKIIα::ChR2 mice, both sexes, 19-28 g) were maintained on a 12 h light/12 h dark cycle with food and water available ad libitum (24–26 °C, 30–40% relative humidity). CaMKIIα::ChR2 mice were obtained by crossing CaMKIIα-Cre mice (Jackson Laboratory; T29-1; Stock No: 005359) with Cre-dependent ChR2(H134R)/EYFP expressing mice (Jackson Laboratory; Ai32; Stock No: 024109). For some pilot optogenetic experiments that require virus injection and optogenetic controls, CaMKIIα-Cre males were used instead. C57BL/6J males (Jackson Laboratory; B6; Stock No: 000664) were used for the initial characterization of the muscimol-induced coma-state. All animal procedures and experimental protocols followed the National Institutes of Health Guide for the Care and Use of Laboratory Animals and were approved by the licensing committee. Animal facilities were reviewed and approved by the institutional animal care and use committee (IACUC) of the University of Wisconsin-Madison and were inspected and accredited by the association for assessment and accreditation of laboratory animal care (AAALAC).

### 3. METHOD DETAILS

#### Surgical procedures

Surgery was performed under isoflurane anesthesia (2.0% induction; 0.8-1.5% maintenance) following aseptic techniques. For muscimol injection, all mice were implanted in the midbrain (A/P −3.75, M/L ±1.00, D/V −1.75) with bilateral guide cannulas (Plastics One). Dummy cannulas were inserted in the guide cannulas to prevent contamination and clogging. To perform electrophysiological recordings, all mice were also implanted with laminar silicon probes (NeuroNexus; A1×16; 50μm spacing), EEG screws, and electromyogram (EMG) wires. One silicon probe was implanted deeply in the right frontal cortex (A/P +1.93, M/L +0.40) to target Area (A)24a (i.e. anterior cingulate cortex, Cg2) and A25 (i.e. infralimbic cortex, IL). For simplicity, we refer to this entire targeted area as mPFC (medial prefrontal cortex) (Carlen, 2017; Laubach et al., 2018). Another silicon probe was implanted in the left posterior parietal cortex (PPtA; A/P −2.00, M/L −2.20) to target all layers. EEG screw electrodes were implanted over left secondary motor cortex (M2; A/P +2.50, M/L −1.50) and right secondary somatosensory cortex (S2; A/P −1.30, M/L +4.0). Reference screws were implanted over the cerebellum and olfactory bulb. EMG stainless steel wires were inserted bilaterally in the dorsal neck musculature and in the whisker musculature. Optic fibers (Doric Lenses; core diameter = 200μm; NA = 0.22; diffuser layer tip) were implanted for optogenetic stimulation. For frontal stimulation, one (n = 4 mice) or two (n = 2 mice) fibers were implanted in the right frontal cortex (A/P +1.77, M/L −0.60) to target mPFC. For parietal stimulation, one (n = 2 mice) or two (n = 6 mice) were placed on the cortical surface over PPtA (A/P −2.00, M/L ±1.80). The craniotomies and silicon probes were covered with surgical silicone adhesive (Kwik-Sil) and all implants were fixed to the skull with dental cement (C&B-Metabond, Fusio or Flow-It). After surgery, mice were individually housed in transparent plastic cages (Allentown Caging; 24.5 x 21.5 x 21cm). At least one week was allowed for recovery from surgery, and experiments started only after the temporal organization of sleep and wakefulness had normalized.

#### Experimental procedures and design

##### Chronic sleep/wake recordings and sleep scoring

After recovering from surgery, mice were connected and accustomed to the recording system, and regularly monitored to ensure that the 24-hour patterns of sleep and waking were normal. Electrophysiological recording and optogenetic stimulation were performed using RZ2 BioAmp processor, OpenEx and Synapse software (Tucker-Davis Technologies). Silicon probes were connected through a head stage to an amplifier (Tucker-Davis Technologies; PZ5 NeuroDigitizer Amplifier) before reaching the RZ2 processor. EEGs and LFPs were filtered by 0.1-100Hz, and multi-unit activities (MUAs) were filtered by 0.3-5kHz. Sampling rate for storage was 256Hz for LFPs, EEGs and EMGs; 25kHz for MUAs. Spike data were collected discreetly from the same LFPs channels. Amplitude thresholds for online spike detection were set manually based on visual control. Whenever the recorded voltage exceeded a predefined threshold, a segment of 46 samples (0.48 ms before, 1.36 ms after the threshold crossing) was extracted and stored for later use. Sleep scoring was performed manually using SleepSign. Analysis of electrophysiological data was performed in MATLAB R2019b and R2021a (MathWorks^®^). LFP data were visually inspected to remove artefacts. Isolated bad channels were replaced by the mean of the immediately surrounding good channels. All LFP channels were subjected to linear detrend and lowpass filtering (200 Hz), using a zero-phase distortion third order Butterworth filter.

##### Induction of a coma-like state

Continuous electrophysiological recordings started at light onset the day before the injection of muscimol and ended the day after the induction of the coma-like state. All sessions were recorded with video. In the afternoon of the experimental day, the first behavioral battery was performed during NREM sleep and wake to obtain baseline levels of responsiveness for each individual mouse (see below). Mice were then briefly anesthetized with sevoflurane (5.0% induction; 3.0% maintenance) to facilitate muscimol injection. After the removal of the dummy cannulas, muscimol (Sigma-Aldrich; M1523; 1mg/mL in saline) was delivered through an internal cannula (Plastics One) whose tip extends beyond the implanted guide cannulas by 2 mm to reach the target area. With a microdialysis pump (Harvard Apparatus, CMA 400) a total of 0.5-0.75 μL of muscimol was injected at a rate of 1 μL/min into the left cannula. After a 30 s pause to limit backflow, the internal cannula was transferred to the right site, and the same procedure was repeated. Anesthesia was discontinued as soon as the procedure was completed. In control experiments (sham injection) the animal was injected with saline according to the same protocol described above for muscimol, then tested behaviorally and monitored until the emergence from anesthesia and recorded until the next day.

##### Behavioral test battery

To assess responsiveness, we used a customized scale developed in our laboratory.

Video and detailed behavioral notes for all mice were collected starting as soon as sevoflurane was discontinued after muscimol injection and lasted until RORR following the coma-like state. The behavioral battery was performed first during NREM sleep and/or wake before the injection (see above), then about 15 min after LORR, and then repeated approximately every 30 to 60 min for the duration of the coma-like state. At least one other behavioral battery was performed shortly after RORR, and a final round was delivered the following day (see Figure 5A for one example in one mouse).

The battery included six sequential tests to assess the response to auditory, tactile, olfactory, vestibular and nociceptive stimuli. The result of each test was scored as 0 (no response), 1 (unclear response) or 2 (positive response) based on the mouse behavior within 5 sec from the onset of the stimulus. The six tests included 1) finger snap at a fixed distance from the mouse cage (0 = no response; 1 = small body tremor; 2 = clear body movement, usually a startle/awakening from sleep); 2) a drop of water was released about 5 cm above the neck of the mouse (0 = no response; 1 = small body tremor; 2 = full movement usually with grooming/awakening from sleep); 3) a freshly opened alcohol swab (BD Alcohol Swabs, 70% v/v Isopropyl Alcohol) was placed about 5 or 1cm in front of the mouse (0 = no response with either distance; 1 = withdrawal/head turning/grooming response only at 1cm distance; 2 = withdrawal/head turning/grooming response at 5 cm distance/awakening from sleep); 4) tail suspension: the mouse was gently picked up by the tail and suspended just above the ground (0 = no response; 1 = some/occasional kicks; 2 = strong body and legs’ movements throughout the test/ awakening from sleep and moves away); 5) righting reflex: the mouse was gently flipped on its side by the tail (0 = laying on its side; 1 = flips back onto its feet after laying on its side; 2 = never really laying on its side/awakening from sleep and moves away); 6) tail pinch (0 = no response; 1 = small movement; 2 = runs away/rights up its body).

##### Optogenetic stimulation

Optic fibers were connected to a blue laser station (473nm, OEM Laser Systems DPSSL Driver, 100mW) controlled by the TDT system, with the laser output power manually regulated by an analog control knob on the driver. Based on the specific excitation threshold of channelrhodopsin ChR2 (Nagel et al., 2005; Madisen et al., 2012; Sidor et al., 2015), and the intended activation radius in the target area, the corresponding laser power was estimated using an established online calculator (https://web.stanford.edu/group/dlab/cgi-bin/graph/chart.php). After the mice were accustomed to the system, and baseline recordings were acquired, brief laser pulses were delivered during NREM sleep to allow a within-mouse calibration of the effective laser power to deliver during the coma-like state. The minimal laser power that induced EEG activation and woke up the mouse from NREM sleep served as a reference for future experiments. To control for possible changes in sleep depth due to time of day, stimulation experiments during sleep, coma, or anesthesia were performed approximately 5 to 8 hours after the beginning of the light phase, when sleep pressure in mice is low (Cavelli et al., 2023). In a few cases the stimulation during sleep was delivered within the first hour of the light phase, and the results appeared to match those obtained when the laser pulses were given later during the day. Overall, across all stimulation experiments during sleep, anesthesia, and the coma-like state, the laser power ranged from 0.2 to 10 mW at fiber tip. Square pulse/s (0.1, 0.25 or 0.5Hz, lasting 1-5 sec) or trains (10ms pulse width; 4, 5, or 8Hz) were delivered. Stimulation during the coma-like state was delivered approximately 30 min after LORR. In a control CaMKIIα-Cre mouse (no virus injection), laser stimulation during NREM sleep, REM sleep or sevo-dex anesthesia did not affect behavior or EEG cortical activity.

##### Anesthesia experiment

At least one week after the induction of the coma-like state another optogenetic stimulation experiment was performed after the mice were deeply anesthetized with sevo-dex (1-2% sevo, 70 or 100ug/kg dex). The dose of dex was 70ug/kg IP in males and 100ug/kg in females, because in our pilot experiments we noticed that female mice required higher doses of dex. Sevo level was then adjusted to reach a state with slow waves in the EEG and stable LORR for at least 5min. The same pattern of stimulation used during the coma-like state, and the same or higher laser power, were used under anesthesia. Electrophysiological and video recordings were performed throughout the duration of the experiment.

##### Histology

Under deep isoflurane anesthesia (3.0%), mice were transcardially perfused with saline (for 30 secs) followed by 4% paraformaldehyde in phosphate buffer. Brains were removed and postfixed for 24 hours in the same fixative, then cut in 50um thick coronal sections on a cryostat (CryoStar^TM^ NX50 or Leica CM1900) after cryoprotection and flash-freezing. Sections were collected in PBS, mounted, air-dried, cover slipped (DAPI-Fluoromount-G, Vectashield, or Permount) and examined under a fluorescent or confocal microscope (Leica, Olympus). In some cases the silicon probe shanks were coated with CM-DiI dye (Thermo Fisher) immediately before implantation for better visualization of the probes’ track. Glial fibrillary acidic protein (GFAP) staining was performed in some mice to localize the silicon probes without fluorescent dye coating (rabbit-anti-GFAP primary antibody, DAKO Z0334, 1:1000 in blocking solution; Donkey-anti-Rabbit AF594 secondary antibody, 1:500 in blocking solution). Crystal Violet staining was also performed in some animals to better visualize the location of the cannulas. EYFP amplification staining was performed in two CaMKIIα::ChR2 mice to confirm the broad cortical expression of the opsin (rabbit anti-GFP primary antibody, Invitrogen, A11122, 1:1000 in blocking solution; goat anti-rabbit Alexa-488 conjugated secondary antibody, Invitrogen, A11008, 1:1000 in blocking solution).

### 4. QUANTIFICATION AND STATISTICAL ANALYSIS

**Statistical Analysis** was performed using linear mixed effect models (Laird and Ware, 1982). The use of mixed effect models allows us to account for repeated measurements (the same mouse being used in multiple experiments). Due to the unbalanced nature of the experiment design, mixed effect models are preferred to traditional repeated measures ANOVA. Parameter estimation of LME models was performed using numerical maximum likelihood estimation, implemented in R by the lmer() function of the lme4 package (Bates et al., 2015). Assumptions were assessed for all models using a scatterplot of fitted values vs. estimate residuals (constant variance) and a QQ-plot of estimated residuals (normality). For all models, the plots indicated normal (or approximately normal) distributions. For the LME models, hypothesis testing is performed by fitting a reduced model with the factor of interest removed, and then comparing the fit of the two models using the asymptotic χ^2^ test. To measure effect size, we took the ratio of the effect of interest (a parameter in the LME model) divided by the combined residual and random effect standard deviation (analogous to Cohen’s D). Below, we report the p-values for each test, additional details (confidence interval, effect size, sample size) are reported in Table 1.

To compare behavioral score in response to stimulation between NREM and REM sleep, as well as different stimulation frequencies, we fit an LME model for behavioral score with state (NREM, REM), position (mPFC, PtA) and stimulation (0.5Hz, 5Hz, 8Hz) as a fixed effects and mouse as a random effect. We found a significant effect of state on the behavioral scores (p = 4.1e-5), and a significant effect of position (p = 0.0002). To compare the mPFC and PtA groups before stimulation, we fit LME models with position (mPFC or PtA) as a fixed effect and mouse as a random effect. We found no significant effect of position when the response variable was the latency to LORR (p = 0.6302) or the behavioral score after LORR (p = 0.8612). To compare behavioral score between NREM sleep and muscimol-induced coma, we fit a LME model for behavioral score with state (NREM vs. pre-stimulation coma) as a fixed effect and mouse as a random effect. We found a significant effect of state on the behavioral scores (p = 2.9e-13).

For stimulation strength, we fit an LME model for strength (frequency + duration) with position (mPFC, PtA), state (REM, NREM), and frequency (0.5 Hz, 5 Hz, 8 Hz) as fixed effects, and mouse as a random effect. We found a significant effect of both position (p = 0.0001) and state (p = 0.00003).

